# Endoglin regulates the integrity of the bone marrow vasculature

**DOI:** 10.1101/2024.07.24.605039

**Authors:** Diego Rodriguez, Mangesh Jaykar, Deepika Watts, Anupam Sinha, Diana Gaete, Anja Krüger, Peter Mirtschink, Martina Rauner, Triantafyllos Chavakis, Helen M. Arthur, Ben Wielockx

## Abstract

Endoglin (Eng) is an accessory receptor for transforming growth factor-β (TGF-β) that is critical for maintaining vascular integrity. Mutations in Eng cause hereditary hemorrhagic telangiectasia (HHT), resulting in arteriovenous malformations (AVMs) and blood abnormalities. Despite the known association between Eng deficiency and AVMs, the underlying mechanisms are unclear. In addition, the role of the bone marrow (BM), a major source of immune and blood cells, in endothelial Eng (EC-Eng) deficiency is unexplored. We show that BM blood vessels conditionally deficient in Eng (cKO) undergo a structured remodeling process over four weeks, with distinct proliferative and resolution phases. These phases involve angiogenic set points, the involvement of integrins, and the modulation of vascular integrity. In addition, we observe changes in hematopoietic stem and progenitor cells (HSPC) and circulating granulocytes, along with reduced red blood cells and platelets due to splenic sequestration. Using a conditional heterozygous EC-Eng deficient mouse model, reflecting the genetics of HHT patients, we identify vascular changes similar to those in the cKO model. Taken together, using multiple *in vivo* approaches, we suggest that reduced Eng expression in the endothelium drives significant BM vascular remodeling, sharing mechanisms with early vascular processes associated with AVM formation.

**Explanation of Novelty:** Our findings reveal that BM blood vessels deficient in endoglin undergo an orchestrated remodeling process with distinct proliferative and resolution phases over several weeks. We identify specific angiogenic set points and profound alterations in vascular integrity, along with hematopoietic changes starting at the level of hematopoietic stem and progenitor cells. These findings advance our understanding of the role of Eng in vascular remodeling and may provide novel therapeutic targets for HHT.

**Key Points:**

1. Conditional EC-Eng deficiency leads to vascular remodeling in the BM of mice in a temporally orchestrated manner.
2. EC-Eng facilitates vascular integrity, hematopoietic homeostasis, and immune cell mobilization.

## Introduction

Endoglin (Eng or CD105) is a transmembrane glycoprotein predominantly expressed in endothelial cells (ECs) and plays a pivotal role as an accessory receptor for members of the Transforming Growth Factor-Beta (TGF-β) family, which is essential for the regulation of cellular functions such as proliferation, differentiation, and migration. It has also been implicated in vascular remodeling in pathological conditions including cancer and cardiovascular disease, highlighting its importance in both normal and diseased states.^1,2^ Notably, endoglin expression is essentially regulated by hypoxia signaling, specifically by the Hypoxia Inducible Factor-1 (HIF-1), highlighting the complexity of cellular responses to environmental stress and its significant impact on vascular dynamics and pathology.^3^ Research efforts have extensively investigated the impact of aberrant endoglin expression on vascular health, revealing that reduced levels of endoglin disrupt endothelial cell physiology and impair angiogenesis, leading to defective endothelial architecture. Furthermore, loss of function mutations in the endoglin gene cause the disease hereditary hemorrhagic telangiectasia (HHT) type 1. This autosomal dominant disorder, characterized by abnormal blood vessel formation, illustrates how mutations in endoglin result in clinical manifestations such as epistaxis and arteriovenous malformations (AVMs) in critical organs, posing serious health risks.^4–9^ Although virtually all endothelial cells in HHT patients suffer from haploinsufficiency, AVMs are rare, suggesting the occurrence of “additional hits” for their manifestation, such as somatic mutation, mechanical trauma and/or inflammation.^10–12^ This complexity is further illustrated by studies correlating AVMs with decreased endoglin expression and abnormal responses to angiogenic stimuli such as VEGF.^13,14^ The angiogenic process itself is a critical aspect of vascular remodeling and health, relying on the orchestrated interaction of various cellular and molecular players, including endoglin.^15^ Its precise initiation, elongation, and termination require the interaction of various cellular components and molecular pathways within ECs and their niche. Interestingly, HHT affects not only vascular integrity but also the immune system, with patients exhibiting increased susceptibility to infection, suggesting a broader systemic impact of endoglin dysfunction.^16,17^ Given that the bone marrow (BM) is a highly vascularized and highly regulated environment for the entire immune system, understanding endoglin signaling in the BM niche is critical for both vascular and hematopoietic health.^18^ For example, aberrant angiogenesis and endoglin can have effects on endothelial function, including induction of permeability, which affects white blood cell (WBC) transmigration.^15,19,20^ The bone marrow vasculature is therefore an important focus for studying how endoglin insufficiency might affect hematopoiesis, including the dynamics of immune cells and hematopoietic progenitors.

In the current study, the role of endothelial endoglin in BM vascular remodeling and its impact on hematopoiesis and peripheral blood dynamics was investigated. Using a conditional knockout mouse model of *Eng* in ECs (Cdh5:CreERT2-*Eng*^f/f^), we analyzed in detail the temporal changes during BM vascular remodeling and the resulting hematopoietic cell dynamics. Leveraging this knowledge, we also studied the HHT patient-relevant heterozygous conditional knockout mice and found a clear resemblance to cKO phenotypes, which may shed new light on the early mechanisms leading to AVM appearance in these patients and beyond. Taken together, our approach not only advances our understanding of the role of EC-Eng in the vascular and hematopoietic systems but may also open new avenues for exploring therapeutic strategies in the treatment of HHT and related vascular anomalies, possibly in combination with existing treatments.

## Materials and methods

All information can be found in the data supplement available with the submission of this article.

## Results

### Endothelial endoglin deficiency leads to remodeling of the BM vasculature

Endoglin is predominantly expressed on the surface of ECs where it helps to maintain the vascular architecture. However, its role on the BM vasculature and consequent impact on hematopoiesis and blood cell dynamics remains unknown. Therefore, we conditionally knocked out endoglin in adult Cdh5:CreERT2-*Eng*^f/f^ mice (henceforth denoted as cKO) through Tamoxifen administration (Supplementary Figure 1A) resulting in loss of endothelial Eng in the BM from one week after the first injection (Figure 1A-B). Next, we analyzed mice at various intervals measuring vessel areas across individual samples (150-400 vessels per mouse) (Figure 1C) and observed a time-dependent remodeling of the vasculature, with a significant reduction in ‘vessel area mode’ (Supplementary Figure 1B) from week three and stabilizing by week four (confirmed up to week nine) (Figure 1C and Supplementary Figure 1C). Despite these changes in vessel area, the total blood vessel area remained largely consistent (Supplementary Figure 1D). To gain insight into the dynamics of changes in vascular integrity over time, we assessed the permeability of the BM vasculature. This was achieved by measuring the extent to which intravenously injected Texas Red-labeled dextran (70kDa) could be detected in the BM, having leaked out of the blood vessel. Consistent with the observed changes in vessel area, we detected more dextran in the BM adjacent to endomucin-positive blood vessels from week three (Figure 1D-E and Supplementary Figure 1E), suggesting that the remodeling of the BM vasculature due to loss of EC-Eng induced leakiness.

**Figure 1.**
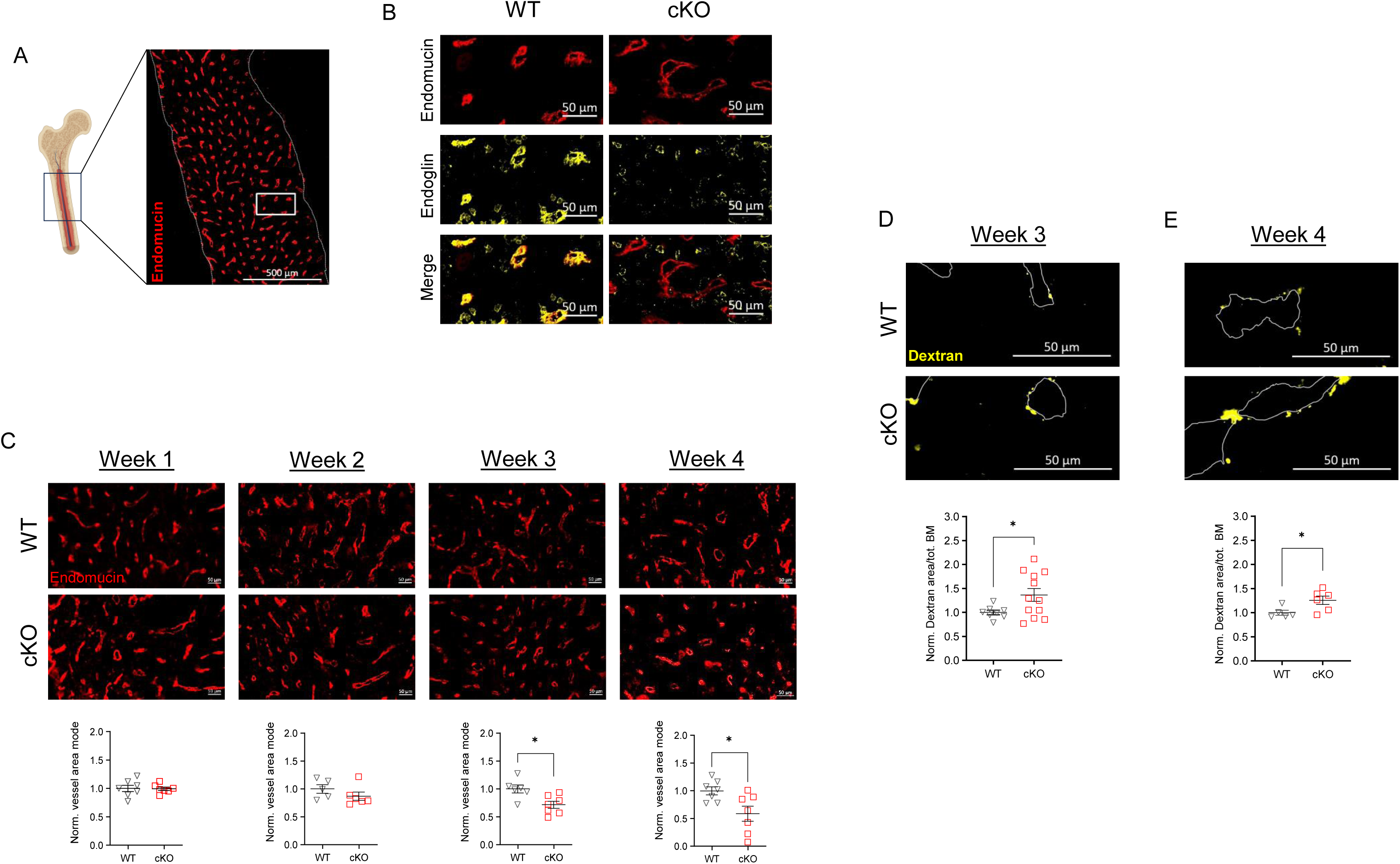
Induced Eng-deficiency in EC results in reduction of blood vessel area and leakage in BM. (A) Representative immunofluorescent (IF) image of endomucin (EC) in the BM. The white insert includes the WT area represented in B. (B) Representative IF images of endomucin and/or Eng in the BM of WT mice and Cdh5:CreERT2-*Eng*^f/f^ littermates one week after initiation of tamoxifen treatment. (C) Representative Endomucin IF in BM sections at one, two, three and four 4 weeks after initiation of tamoxifen treatment and corresponding normalized vessel area modes (µm^2^). (D-E) Representative IF images of dextran-Texas Red (yellow) leaked into the BM (note: adjacent vessels are represented by dotted lines) three and four weeks after initiation of tamoxifen treatment; including the normalized dextran^+^ area in the total BM (%).

### EC-Eng deficiency affects mature hematopoietic cell dynamics in a time-dependent manner

Next, we studied whether these changes affected immune cell dynamics in the circulation. Initially, monocyte and B-cell levels increased in the third week, returning to baseline by the fourth week (Figure 2A and Supplementary Figure 2A), whereas neutrophils and eosinophils exhibited a similar increase at week 3 but remained significantly elevated thereafter (Figure 2B and Supplementary Figure 2A). Contrarily, RBC and PLT counts decreased early and persisted below WT control levels (Figure 2C and Supplementary Figure 2B). However, the increase in their circulating precursors, reticulocytes and reticulated platelets, respectively, suggests their increased production in the BM to compensate for these losses (Supplementary Figure 2C-D). This observation led us to investigate the spleen, recognizing its critical role as a principal filter of the blood. The spleen is capable of sequestering circulating RBCs and PLTs, in addition to its function in expelling senescent or modified RBCs through the action of resident red pulp macrophages (RPM).^21^ Here, we reveal a significant increase in the cKO spleen weight starting early after cre-recombinase activation (Supplementary Figure 2E). This was accompanied by a substantial rise in the count of RPM per spleen and an expansion of CD42b-positive cells (active PLT) (Supplementary Figure 2F-G). However, we detected no differences in pro- and anti-inflammatory cytokines in circulation throughout the first four weeks, indicating that these cellular changes were not linked to a profound systemic inflammation (Supplementary Figure 3A). Taken together, these results strongly suggest that the BM enhances the production of both RBC and PLT as a compensatory response to their accelerated clearance by the spleen, while the increased leakiness of the BM vasculature significantly affects mature immune cell dynamics.

**Figure 2.**
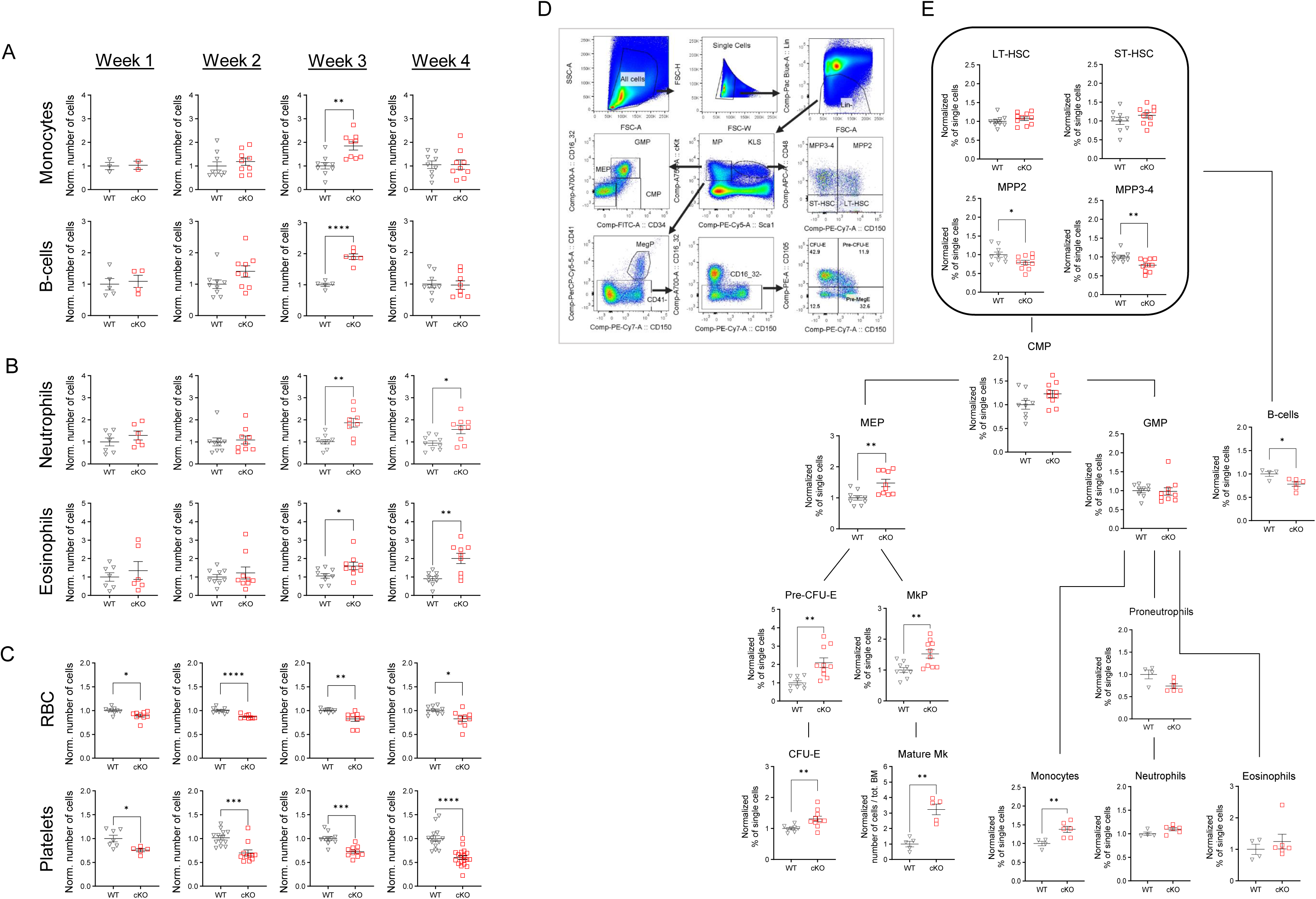
Induced Eng-deficiency in EC affects hematopoietic cell dynamics in a time dependent manner. (A-C) Normalized number of different blood cells in circulation from WT mice and Cdh5:CreERT2-*Eng*^f/f^ littermates at different time points after treatment with tamoxifen (#cells/µl). (D) FACS gating strategy for identification of HSPC by staining surface markers as detailed in the Methods section (note: no differences in CD105 levels were detected for any of the hematopoietic cell lineages between both genotypes) (E) Normalized percentage of single cells in the bone marrow of WT mice and Cdh5:CreERT2-*Eng*^f/f^ littermates four weeks after initiation of tamoxifen treatment. Each graph represents data from three independent experiments.

### Re-organization of the hematopoietic stem and progenitor cells (HSPC) in the bone marrow as a direct consequence of loss of EC-Eng

Next, we focused on the BM HSPCs to better understand how the previously described changes in the vasculature might affect the hematopoietic framework, using a comprehensive FACS analysis (Figure 2D). Οur results reveal that at the pivotal fourth week multipotent progenitor (MPP) segments from EC-Eng deficient mice are significantly decreased as compared to WT littermates (Figure 2E), although no significant differences were observed in the populations of common myeloid progenitors (CMP), granulocyte-monocyte progenitors (GMP) (Figure 2E). Even though the initial increase in the number of circulating monocytes and B-cells had normalized by week four, the number of mature monocytes significantly increased while B-cells significantly decreased. This observation aligns with prior findings that delineated a distinct disparity in the production rates of monocytes compared to B lymphocytes.^22,23^ In contrast, progenitors responsible for RBC and PLT production, including Megakaryocyte-Erythrocyte-Progenitors (MEP), Pre-Erythroid Colony-Forming Unit (Pre-CFUE), Erythroid Colony-Forming Unit (CFUE), Megakaryocyte Progenitors (MkP) and mature megakaryocytes (Mk) exhibited a marked increase, which underscores the enhanced production of reticulocytes and reticulated platelets in circulation (Figure 2E and Supplementary Figure 2C-D). Moreover, although vascular remodeling in the BM has been linked to bone formation,^24,25^ loss of EC-Eng did not alter the integrity of the bone (Supplementary Figure 3B). Taken together, these findings demonstrate the wide impact of EC-Eng on the entire hematopoietic system in the BM and beyond (Supplementary Figure 3C).

### Induced loss of EC-Eng causes vessel remodeling through endothelial cell proliferation and subsequent apoptosis

To capture the nuances of the temporal impact of endoglin deficiency, we next evaluated the proliferative behavior and survival rates of BM EC (Figure 3A). Comprehensive FACS analyses revealed a significant decrease in Ki67-negative cells (non-proliferative or quiescent cells - G_0_ phase) at weeks two and three, indicating increased cell cycle activity. This enhanced proliferation returned to WT-like levels by week four (Figure 3B). In parallel, the proportion of viable ECs, as assessed by Annexin V and/or DAPI, remained stable for the first three weeks but showed a significant reduction in Eng-deficient ECs by week four (Figure 3C). Notably, the total number of endothelial cells within the bone marrow was consistent throughout the four-week period (Figure 3D), mirroring the findings shown in Supplementary Figure 1D. This intricate sequence of events supports the concept that Eng deficiency precipitates a two-phase vascular remodeling process within the BM. Concurrently, the pronounced fluctuation in white blood cell counts during this period (Figure 2A-B), suggests a transient destabilization of vascular integrity, particularly during the third week.

**Figure 3.**
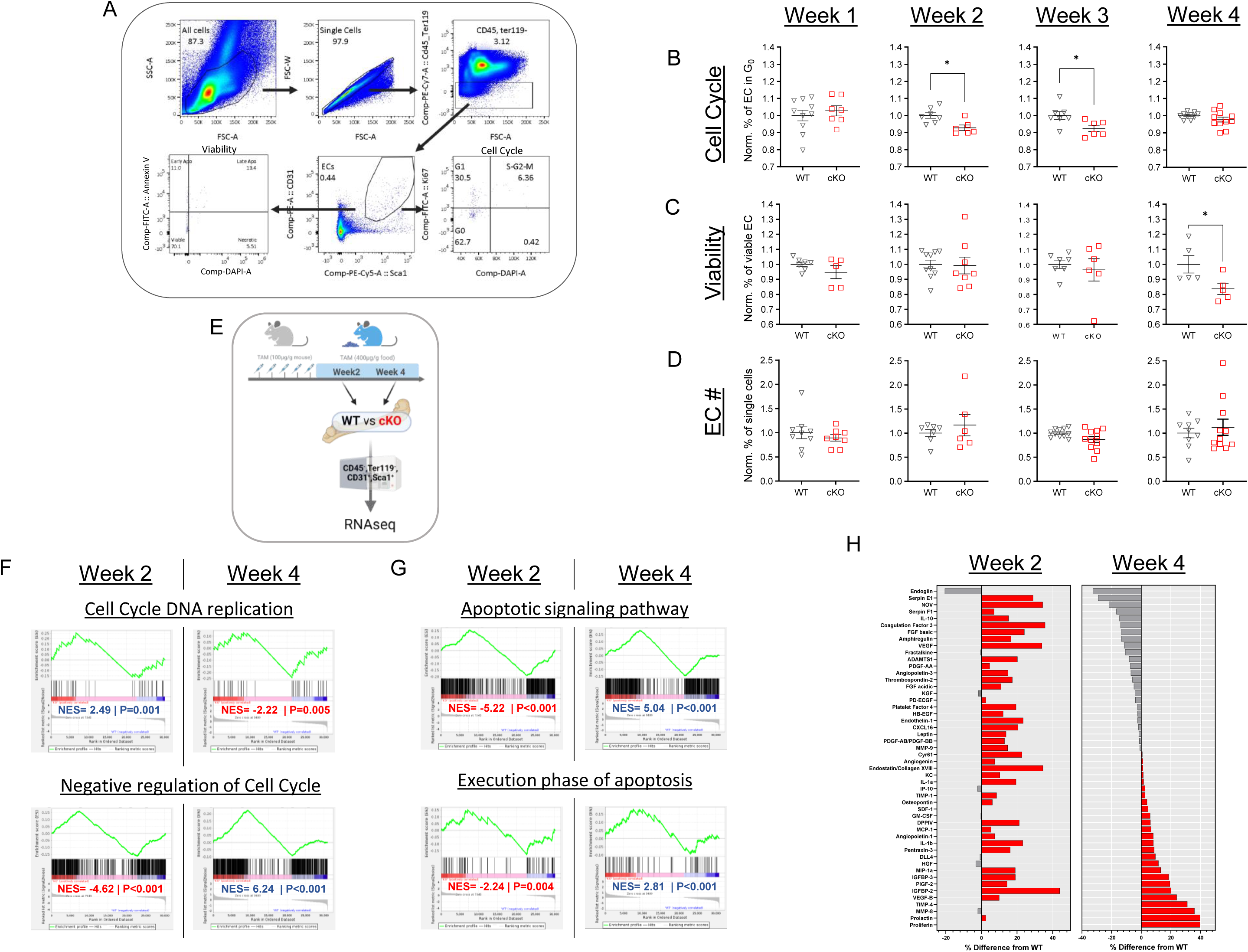
Loss of Eng in the BM-EC leads to remodeling of the vasculature in a temporally and orchestrated manner. (A) FACS gating strategy to identify EC, their viability and cell cycle status as described in the Methods section. (B-C) Normalized percentage of quiescent (G_0_) and viable EC in the BM (%). Each graph represents data from two to three independent experiments. (D) Normalized number of EC as percentage of total number of cells (%). Each graph represents data from two to three independent experiments. (E) Experimental design to obtain RNAseq data from sorted BM-EC from the different genotypes. (F-G) Gene set enrichment analysis (GSEA) for samples at two weeks and four weeks after the start of tamoxifen treatment. (H) Mouse angiogenesis array proteome profiler using BM extracellular protein content as described in the Methods section. Each bar represents the mean of two independent samples and each sample was a pool of three different mice. Data is shown as the percent difference of the WT average.

To deepen our understanding of the temporal nuances in EC behavior, we performed a comprehensive analysis by bulk RNA sequencing of sorted BM ECs (characterized as CD45^-^, Ter119^-^, CD31^+^, Sca1^+^) from both WT and cKO mice at the critical junctures of week two and week four (Figure 3E). As expected, gene set enrichment analysis (GSEA) revealed a pronounced decrease in the activation of TGFβ and BMP pathways in cKO ECs at both time points (Supplementary Figure 4A). This observation is consistent with the established function of endoglin as an auxiliary receptor for TGFβ^26^ and its high affinity for BMP9 and -10.^27^ Consistent with our previous observations, we detected marked differences in cell cycle activity between the two time points. Specifically, there was an increase in cell cycle-related gene signature at week two (Figure 3F), indicative of increased endothelial proliferation, which included increased VEGFR2 binding activity, a key regulator of angiogenesis (Supplementary Figure 4B).^28^ Conversely, week four was characterized by a significant increase in apoptosis-related gene signatures, indicating a shift towards endothelial cell resolution (Figure 3G).

To assess the impact of this remodeling phase, we evaluated the extracellular protein landscape within the BM at weeks two and four using a mouse angiogenesis array proteome profiler. The findings at week two contrasted sharply with those at week four, with widespread overexpression of angiogenesis related proteins in cKO mice (Figure 3H). However, by week four, the expression profile had converted to a more balanced state between the two genotypes (Figure 3H). Together, these findings elucidate the orchestrated temporal progression of BM vasculature remodeling following Eng deficiency, modulating the complex interplay within the bone marrow environment.

### Eng deficiency reduces endothelial adhesion and modulates integrin dynamics in a time-dependent manner

The differences between week two and week four indicate a highly orchestrated and complex process that finally results in a new balance associated with a permanent increased permeability. Therefore, we searched for additional differences during the different time points and found great fluctuations in endothelial barrier properties including “homotypic cell-cell adhesion” and “tight junctions” (Figure 4A). Furthermore, in our analysis of EC behavior and vascular remodeling within the BM, we observed a profound downregulation in “cell-matrix adhesion” signature at both week two and week four (Supplementary Figure 5A). This might appear contradictory, given that increased vascular permeability is observed only from week three. However, these findings underscore the dynamic and multifactorial nature of endoglin-deficiency in endothelial remodeling, suggesting that while downregulation of cell matrix adhesion is a consistent feature across the observed timeframe, the impact on vascular integrity and leakiness is modulated by additional changes. To further elucidate these dynamic mechanisms, we applied Ingenuity Pathway Analysis (IPA) to our RNAseq data. Interestingly, we discovered a significant shift in the activity profiles of β-integrins across the critical intervals of week two and week four. Specifically, the activity of virtually all β-integrins was predicted to be increased at week two in cKO ECs but significantly decreased at week four, with the exception of integrin beta 5 (ITGB5) (Figure 4B).

**Figure 4.**
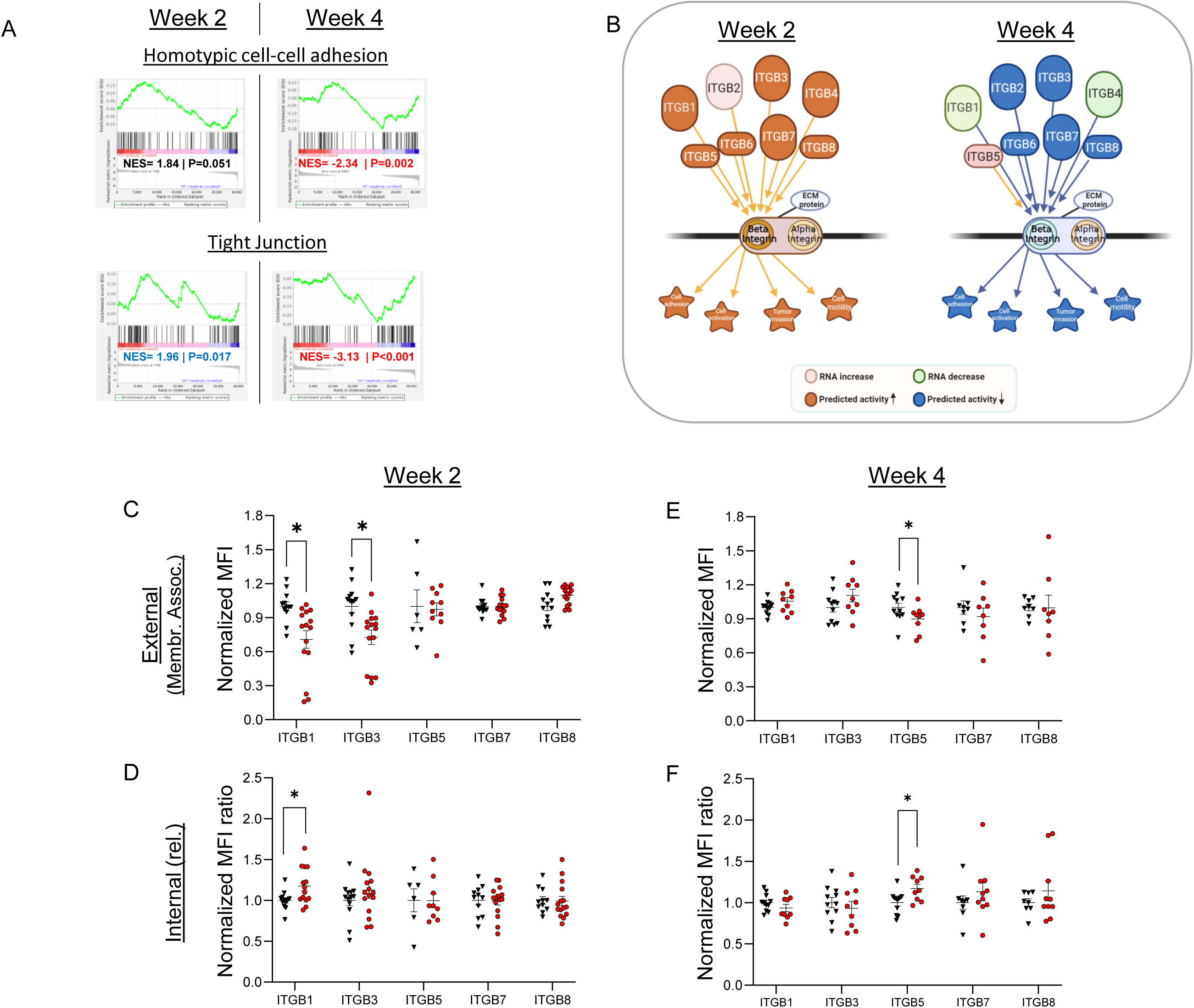
Time dependent modulation of BM endothelial adhesion and integrin dynamics. (A) GSEA displays differential changes in homotypic cell-cell adhesion and tight junction gene signatures. (B) Ingenuity Pathway Analysis (IPA) predicts that the activity of β-integrin family members in EC from cKO mice at week two is drastically different from the predicted activity at week four. (C, E) Median fluorescence intensity (MFI) of 5 ITGβ family members on non-permeabilized e-selectin^+^ EC (=external). (D, F) The ratio of MFI of ITGβ family members on permeabilized versus non-permeabilized e-selectin^+^ EC (=Normalized fraction). Each graph (C-F) represents data from two to three independent experiments.

Next, we investigated the physical response of β-integrins to the switch from proliferation to apoptosis. For this, we analyzed their localization on and within ECs, as this could coincide with the activity of pro-angiogenic factors known to induce β-integrin endocytosis. Using flow cytometry, we measured the mean fluorescence intensity (MFI) of β-integrins on the surface of activated (E-selectin^+^) ECs (Figure 4C & E) and after permeabilization of the cells (total β-integrins= surface + internal) (Supplementary Figure 5B-C). Quantification of the total β-integrins versus surface ratio provided insight into their dynamic intracellular distribution over time (Figure 4D & F). Taken together, we observed a reduction in surface ITGB1 and 3 at week two (Figure 4C), associated with a significant increase in intracellular ITGB1 and a positive trend for ITGB3 (Figure 4D). Interestingly, ITGB5 followed a similar pattern to ITGB1, but only at week four, as predicted by IPA (Figure 4C). Despite these shifts, the total amount of integrins remained consistent across time points (Supplementary Figure 5B-C). Taken together, these findings suggest that ITGB1 and possibly ITGB3, transiently undergo increased internalization at week 2, potentially affecting the early phase of the angiogenesis process, ^29–31^ whereas ITGB5 appears to play a role in the later phase of vascular remodeling and maintenance of vascular integrity in the BM.

### Heterozygous Eng-EC deficiency leads to the development of smaller Eng^-^ vessels in the BM

Given that HHT patients carry heterozygous Eng mutations ultimately leading to vascular anomalies in various organs,^17^ we next explored the effects of heterozygous Eng-loss in BM-ECs using the Cdh5:CreERT2-*Eng*^f/+^ mouse model (referred to as Het) and their control littermates. We first analyzed Eng expression in BM-ECs four weeks after tamoxifen treatment using FACS with cKO mice as a negative control (Figure 5A and Supplementary Figure 6A). Surprisingly, this revealed a significant increase in the prevalence of Eng-negative ECs (Eng^-^ EC) in Het mice compared to their WT counterparts (Figure 5B). Next, we examined the spatial distribution of Eng expression within the individual BM vessels. By staining for endomucin and Eng, we observed that Het mice had a greater number of vessels that were either partially (=Eng^part^) or completely endoglin negative (=Eng^-^) compared to their WT littermates (Figure 5C-E). Notably, vessels with no Eng^+^ ECs were uniformly smaller than those showing any Eng^+^ EC (Eng^+^ or Eng^part^) (Figure 5F), which is consistent with the vascular changes previously described in cKO vessels (Figure 1D). This evidence clearly demonstrates that Het mice contain a significantly greater number of Eng^-^ vessels, a finding that has previously been linked to AVM formation.^13^ As the latter has been associated with dramatic widening of vessels, we next quantified all vessels larger than 1500µm^2^. This showed that, although the difference was not significant, Het mice tended to have more very large vessels in their BM. (Supplementary Figure 6B).

**Figure 5.**
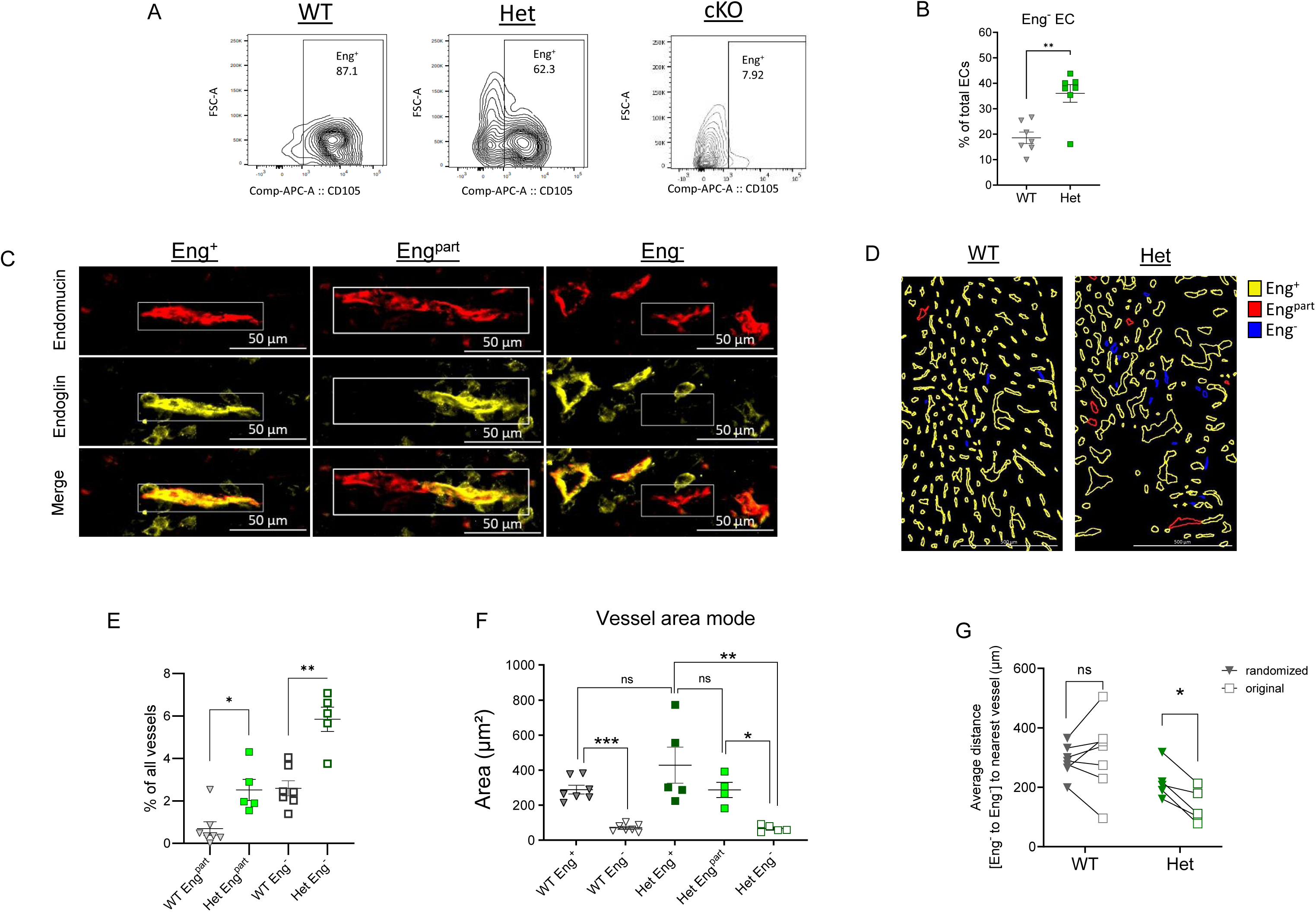
Heterozygous EC-Eng deficiency enhances non-randomly distributed Eng-negative vessels in the BM. (A) Representative FACS plots of Eng-expression in BM-ECs from WT, Hetero and cKO mice. (B) Number of Eng^-^ EC as a percentage of total EC in the BM. (C) Representative IF images of endomucin and/or Eng in the BM of WT mice and Cdh5:CreERT2-Eng^f/+^ littermates four weeks after initiation of tamoxifen treatment. (D) Representative images of the distribution of Eng^+^ (yellow), Eng^part^ (red), and Eng^-^ (blue) region of interest (ROI) in BM sections. (E) Normalized amount of Eng^part^ and Eng^-^ vessels. (F) Vessel area mode of Eng^+^, Eng^part^, and Eng^-^ vessels in Cdh5:CreERT2-*Eng*^f/+^ mice and Eng^+^ and Eng^-^ vessels in WT littermates. (G) Average distance of each Eng^-^ vessel to the nearest Eng^-^ vessel after randomization of all vessels, compared to the distance of the same Eng^-^ vessels in the original situation as displayed in (D).

Finally, we also analyzed the potential association of all smaller Eng^-^ vessels across the BM (Figure 5D), using nearest neighbor analyses as described in Materials and Methods. Interestingly, while the vessels in WT mice were not closer to each other than expected by chance, our data clearly show a closer alignment of Eng^-^ vessels in the BM of Het mice (Figure 5D and G). Taken together, these results suggest that heterozygous deletion of endoglin in BM-EC can lead to a significantly higher level of Eng^-^ ECs closely aligned in smaller Eng^-^ vessels (Figure 1C).

### Heterozygous Eng-deficiency affects immune cell dynamics in a time dependent manner

To determine whether the observed changes would result in impaired vessel functionality, we analyzed BM leakage in Het mice four weeks after TAM treatment using the dextran approach (Figure 6A). Interestingly, Het mice exhibited a significantly larger dextran^+^ area compared to their WT counterparts (Figure 6B), indicating increased vascular permeability, a phenomenon also observed in cKO mice (Figure 1D-E). In addition, we analyzed blood cells and their progenitors over four weeks but observed no gross changes in circulating monocytes, B cells (Supplementary Figure 7A-B), or erythroid and platelet-associated lineages (Supplementary Figure 8A-B). Interestingly, however, and despite the stability of hematopoietic progenitors at week 4 (Supplementary Figure 8C-D), we again detected a significant increase in neutrophils and eosinophils in the blood of Het mice compared to their WT counterparts (Figure 6C-D). This pattern not only mirrors our findings in cKO mice (Figure 1E and 2B), but also underscores the link between Het and cKO mice in terms of Eng^-^ EC/vessels, highlighting a common pathway affecting vascular remodeling and consequent immune cell recruitment.

**Figure 6.**
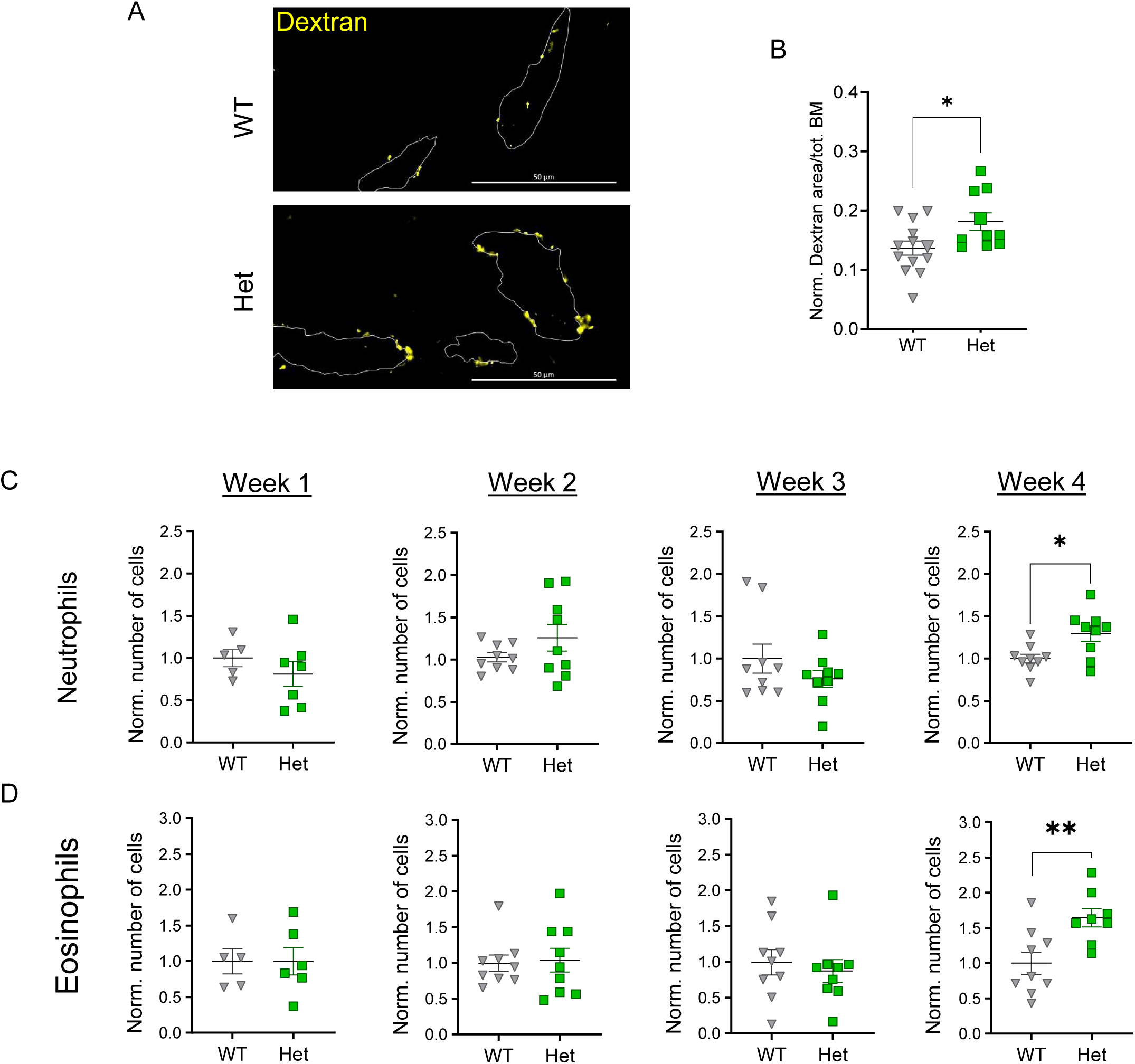
Heterozygous EC-CD105 deficiency leads to increased, non-randomly distributed appearance of CD105 negative vessels in the bone marrow. (A) Representative IF images of dextran-Texas Red leaked into the BM (note: adjacent vessels are represented by dotted lines) four weeks after initiation of tamoxifen treatment. (B) Normalized Dextran^+^ staining per total area of BM (%). (C-D) Normalized number of different blood cells in circulation from WT mice and Cdh5:CreERT2-Eng^f/+^ littermates at different time points after treatment with tamoxifen (#cells/µl).

## Discussion

The aim of this study was to investigate the role of EC-Eng in the regulation of vascular function in the BM environment. Our results show that conditional endoglin deficiency initiates a time-dependent vascular remodeling process characterized by distinct phases of cell proliferation and apoptosis. This remodeling is influenced by a variety of angiogenic factors, including several beta integrins, and ultimately leads to the formation of smaller but more leaky vessels. Notably, our results also show that heterozygous mice (Het), which retain a functional endoglin allele in their endothelial cells, have a significant increase in vessels in which endoglin is not detectable. The area of these vessels was significantly smaller, exhibiting comparable outcomes to those observed in cKO mice. Taken together, our research provides a comprehensive analysis of the temporal dynamics of vascular remodeling in the bone marrow following loss of EC-Eng. Using a combination of in vivo mouse models, deep sequencing, microscopy and flow cytometry, our findings highlight the potential implications of these processes in the development of arteriovenous malformations in HHT patients.

Recent studies have highlighted the critical role of ENG signaling in regulating directional migration and hyperproliferation of ECs, linking defects in this pathway to the development of AVM ^7,32^. In addition, endothelial Eng has been shown to be involved in inflammation, playing an important integrin-mediated role in leukocyte adhesion and vascular transmigration,^33–35^ suggesting that HHT patients may also experience impaired or delayed immune responses. Endothelial integrin activity occupies a fundamental position as it can regulate angiogenesis, vascular integrity, and leukocyte activity.^29,36^ Integrins and TGF-β largely influence each other’s functions,^37^ suggesting that endothelial integrin activity in HHT patients may influence immune responses and AVM development. Our combined results suggest that EC-Eng deficiency affects different β integrins in a time-dependent manner. Specifically, our results suggest that there is a decrease in integrin adhesion at two weeks, possibly mediated by a decrease in the amount of membrane-associated ITGB1 and 3.

ITGB1 and 3 can regulate a range of endothelial activities including barrier integrity ^38^, interaction with leukocytes, PLT, and the extracellular matrix, migration, cell cycle,^36,39,40^ and apoptosis^41^, making them key endothelial regulators. This makes them potential regulators of the vascular permeability observed in the BM following EC-Eng deficiency. In addition, for ITGB5, previous studies suggest opposing activities with respect to endothelial barrier integrity, consistent with the differences observed in FACS and IPA between ITGB1 and 3, and 5, further supporting the idea that these beta integrins are central in the identified differences in cKO mice.

Given the well-established importance of the BM and its vascular niche^42^ in the mobilization of immune cells and maintenance of proper hematopoiesis, we investigated the extent to which conditional reduction of EC-Eng in the BM of adult mice could directly facilitate vascular niche remodeling and affect the hematopoietic system. Furthermore, we examined different time points over a period of four weeks following induction of Eng deficiency. This gave us the exceptional advantage of being able to precisely observe the time course and specific stages at which BM vasculature remodeling occurs in unperturbed adult mice, and led to the observation that remodeling of the Eng-deficient vasculature takes approximately four weeks to complete and is characterized by a proliferative phase during weeks two and three, followed by a resolution phase at week four. To the best of our knowledge, this is the first time that such a detailed temporal analysis has been performed in vivo, providing a comprehensive understanding of the remodeling process associated with Eng loss. Notably, our observation that BM blood vessels became much smaller than those in WT mice is previously undescribed. Previous research suggests that the absence of endoglin typically results in vascular enlargement, a feature also seen in HHT patients.^19^ However, similar to HHT patients, our cKO mice exhibited more fragile and leaky vessels in their BM, as evidenced by elevated levels of circulating granulocytes, even weeks after stabilization of new vascular structures.

Our observation that BM vessels undergo this typical remodeling following loss of EC-Eng might help to better understand the phenotypes seen in HHT patients. In this regard, we found that mice with one remaining functional Eng allele in their ECs (Het mice) have approximately double the number of Eng-negative ECs and display significantly increased numbers of mosaic (Eng^part^) and fully Eng-deficient (Eng^-^) vessels. In addition, the latter vessels were as small as those in cKO mice four weeks after TAM and were more interconnected than expected by chance. Although we could not conclusively determine whether these specific mosaic or Eng^-^ vessels caused lesions, we were able to correlate our histologic findings with increased vascular leakage and higher circulating granulocyte levels. ^13,14^ Taken together, our data clearly indicate that Eng expression at very low to undetectable levels leads to vascular remodeling that eventually reaches a new equilibrium in the BM, even in Het mice. Notably, the incidence of AVMs appearance in patients increases with age,^43,44^ suggesting the possibility that a barrage of mild challenges accumulates throughout a vessel’s lifetime, which may be sufficient to culminate in the emergence of an AVM. In this sense, the mechanisms described here, showing decreased Eng expression in vessels that were previously rendered vulnerable by a heterozygous mutation in Eng, may represent the first steps that are necessary but not sufficient for AVM formation. In this case, the true impact of heterozygous Eng deficiency in BM ECs and its effect on the neighboring vasculature remains unclear, and therefore future research is warranted to determine whether Eng^part^ or Eng^-^ vessels can ultimately form AVMs in the BM or whether they represent an intermediate step toward AVM formation.

AVM appearance and associated hemorrhage often lead to hemodynamic abnormalities in HHT patients, including anemia,^45^ which may mask underlying hemodynamic abnormalities. Here, we demonstrated that induced Eng deficiency in cKO mice resulted in time-dependent vascular permeability and a significant early decrease in circulating erythrocytes and PLT. Interestingly, our data suggest that this decrease was directly associated with a dramatic increase in sequestration of both cell types by the more receptive spleen. This ultimately led to a compensatory overproduction of RBCs and PLTs, provoking a complete overhaul of the progenitor populations from the erythroid/megakaryocytic lineages to the multipotent progenitors. In contrast, no such dramatic changes were detected in Het mice four weeks after heterozygous loss of EC-Eng, indicating that the spleen was not as affected.

In conclusion, our results demonstrate that bone marrow EC-Eng deficiency induces vascular remodeling that takes four weeks to complete and involves temporal changes in apoptotic, proliferative, and integrin activity in endothelial cells. This remodeling resulted in increased vascular permeability beginning at week three, which correlated with increased circulating granulocyte counts. The effects of EC-Eng deficiency on the vasculature cascaded to a complete alteration of hematopoietic progenitor populations in the bone marrow and beyond. Importantly, our findings suggest that the mechanisms observed in cKO mice may be relevant to Eng-deficient vessels in Het mice, potentially providing new insights into AVM formation and the interaction between EC-Eng deficiency and blood cells in HHT patients.

## Acknowledgments

This work was supported by grants from the DFG (German Research Foundation) within the CRC/TRR 205/2 (project A02) and CRC/TRR 369/1 (project A01) to B.W. and CRC/TRR 369/1 (project C03) to T.C.; DFG grants WI3291/12-1, 13-1 and 14-3 to B.W. and FOR5146 (to M.R.), and a grant from the priority program µBONE 2084 (to M.R. and B.W.). We thank Ralf Adams (Münster, Germany) for kindly providing the Cdh5:CreERT2 mouse line.

## Authors’ contributions

D.R. designed the study, performed the majority of experiments, analyzed data, and drafted the manuscript. M.J. and D.W. provided technical expertise, analyzed data and contributed to the discussion. A.S. performed bioinformatics analyses. D.G., P.M., A.K. and M.R. provided technical expertise. T.C. and H.M.A. provided essential tools and contributed to data interpretation and discussion, and B.W. designed and supervised the overall study, analyzed data, and wrote the manuscript. All authors discussed the results and commented on the manuscript.

## Notes

https://www.ncbi.nlm.nih.gov/geo/query/acc.cgi?acc=GSE271409

### Competing Interest Statement

The authors have declared no competing interest.

## References

1. Rossi E, Bernabeu C. Novel vascular roles of human endoglin in pathophysiology. J Thromb Haemost. 2023;21(9):2327–2338.

2. Gonzalez Munoz T, Amaral AT, Puerto-Camacho P, Peinado H, de Alava E. Endoglin in the Spotlight to Treat Cancer. Int J Mol Sci. 2021;22(6).

3. Sanchez-Elsner T, Botella LM, Velasco B, Langa C, Bernabeu C. Endoglin expression is regulated by transcriptional cooperation between the hypoxia and transforming growth factor-beta pathways. J Biol Chem. 2002;277(46):43799–43808.

4. Ollauri-Ibanez C, Nunez-Gomez E, Egido-Turrion C, et al. Continuous endoglin (CD105) overexpression disrupts angiogenesis and facilitates tumor cell metastasis. Angiogenesis. 2020;23(2):231–247.

5. Park S, Dimaio TA, Liu W, Wang S, Sorenson CM, Sheibani N. Endoglin regulates the activation and quiescence of endothelium by participating in canonical and non-canonical TGF-beta signaling pathways. J Cell Sci. 2013;126(Pt 6):1392–1405.

6. Jerkic M, Rodriguez-Barbero A, Prieto M, et al. Reduced angiogenic responses in adult Endoglin heterozygous mice. Cardiovasc Res. 2006;69(4):845–854.

7. Mahmoud M, Allinson KR, Zhai Z, et al. Pathogenesis of arteriovenous malformations in the absence of endoglin. Circ Res. 2010;106(8):1425–1433.

8. Sugden WW, Meissner R, Aegerter-Wilmsen T, et al. Endoglin controls blood vessel diameter through endothelial cell shape changes in response to haemodynamic cues. Nat Cell Biol. 2017;19(6):653–665.

9. Faughnan ME, Palda VA, Garcia-Tsao G, et al. International guidelines for the diagnosis and management of hereditary haemorrhagic telangiectasia. J Med Genet. 2011;48(2):73–87.

10. Arthur HM, Roman BL. An update on preclinical models of hereditary haemorrhagic telangiectasia: Insights into disease mechanisms. Front Med (Lausanne). 2022;9:973964.

11. Bernabeu C, Bayrak-Toydemir P, McDonald J, Letarte M. Potential Second-Hits in Hereditary Hemorrhagic Telangiectasia. J Clin Med. 2020;9(11).

12. Snellings DA, Gallione CJ, Clark DS, Vozoris NT, Faughnan ME, Marchuk DA. Somatic Mutations in Vascular Malformations of Hereditary Hemorrhagic Telangiectasia Result in Bi-allelic Loss of ENG or ACVRL1. Am J Hum Genet. 2019;105(5):894–906.

13. Galaris G, Montagne K, Thalgott JH, et al. Thresholds of Endoglin Expression in Endothelial Cells Explains Vascular Etiology in Hereditary Hemorrhagic Telangiectasia Type 1. Int J Mol Sci. 2021;22(16).

14. Al-Samkari H, Kasthuri RS, Parambil JG, et al. An international, multicenter study of intravenous bevacizumab for bleeding in hereditary hemorrhagic telangiectasia: the InHIBIT-Bleed study. Haematologica. 2021;106(8):2161–2169.

15. Rodriguez D, Watts D, Gaete D, Sormendi S, Wielockx B. Hypoxia Pathway Proteins and Their Impact on the Blood Vasculature. Int J Mol Sci. 2021;22(17).

16. Droege F, Thangavelu K, Stuck BA, Stang A, Lang S, Geisthoff U. Life expectancy and comorbidities in patients with hereditary hemorrhagic telangiectasia. Vasc Med. 2018;23(4):377–383.

17. Schoonderwoerd MJA, Goumans MTH, Hawinkels L. Endoglin: Beyond the Endothelium. Biomolecules. 2020;10(2).

18. Moody JL, Singbrant S, Karlsson G, et al. Endoglin is not critical for hematopoietic stem cell engraftment and reconstitution but regulates adult erythroid development. Stem Cells. 2007;25(11):2809–2819.

19. Rossi E, Bernabeu C, Smadja DM. Endoglin as an Adhesion Molecule in Mature and Progenitor Endothelial Cells: A Function Beyond TGF-beta. Front Med (Lausanne). 2019;6:10.

20. Sluiter TJ, van Buul JD, Huveneers S, Quax PHA, de Vries MR. Endothelial Barrier Function and Leukocyte Transmigration in Atherosclerosis. Biomedicines. 2021;9(4).

21. Li H, Liu ZL, Lu L, Buffet P, Karniadakis GE. How the spleen reshapes and retains young and old red blood cells: A computational investigation. PLoS Comput Biol. 2021;17(11):e1009516.

22. Shahaf G, Zisman-Rozen S, Benhamou D, Melamed D, Mehr R. B Cell Development in the Bone Marrow Is Regulated by Homeostatic Feedback Exerted by Mature B Cells. Front Immunol. 2016;7:77.

23. Whitelaw DM. The intravascular lifespan of monocytes. Blood. 1966;28(3):455–464.

24. Kusumbe AP, Ramasamy SK, Adams RH. Coupling of angiogenesis and osteogenesis by a specific vessel subtype in bone. Nature. 2014;507(7492):323–328.

25. Rauner M, Murray M, Thiele S, et al. Epo/EpoR signaling in osteoprogenitor cells is essential for bone homeostasis and Epo-induced bone loss. Bone Res. 2021;9(1):42.

26. Barbara NP, Wrana JL, Letarte M. Endoglin is an accessory protein that interacts with the signaling receptor complex of multiple members of the transforming growth factor-beta superfamily. J Biol Chem. 1999;274(2):584–594.

27. Castonguay R, Werner ED, Matthews RG, et al. Soluble endoglin specifically binds bone morphogenetic proteins 9 and 10 via its orphan domain, inhibits blood vessel formation, and suppresses tumor growth. J Biol Chem. 2011;286(34):30034–30046.

28. Melincovici CS, Bosca AB, Susman S, et al. Vascular endothelial growth factor (VEGF) - key factor in normal and pathological angiogenesis. Rom J Morphol Embryol. 2018;59(2):455–467.

29. Tanjore H, Zeisberg EM, Gerami-Naini B, Kalluri R. Beta1 integrin expression on endothelial cells is required for angiogenesis but not for vasculogenesis. Dev Dyn. 2008;237(1):75–82.

30. Simons M, Gordon E, Claesson-Welsh L. Mechanisms and regulation of endothelial VEGF receptor signalling. Nat Rev Mol Cell Biol. 2016;17(10):611–625.

31. Valdembri D, Caswell PT, Anderson KI, et al. Neuropilin-1/GIPC1 signaling regulates alpha5beta1 integrin traffic and function in endothelial cells. PLoS Biol. 2009;7(1):e25.

32. Jin Y, Muhl L, Burmakin M, et al. Endoglin prevents vascular malformation by regulating flow-induced cell migration and specification through VEGFR2 signalling. Nat Cell Biol. 2017;19(6):639–652.

33. Torsney E, Charlton R, Parums D, Collis M, Arthur HM. Inducible expression of human endoglin during inflammation and wound healing in vivo. Inflamm Res. 2002;51(9):464–470.

34. Rossi E, Lopez-Novoa JM, Bernabeu C. Endoglin involvement in integrin-mediated cell adhesion as a putative pathogenic mechanism in hereditary hemorrhagic telangiectasia type 1 (HHT1). Front Genet. 2014;5:457.

35. Rossi E, Sanz-Rodriguez F, Eleno N, et al. Endothelial endoglin is involved in inflammation: role in leukocyte adhesion and transmigration. Blood. 2013;121(2):403–415.

36. Yamamoto H, Ehling M, Kato K, et al. Integrin beta1 controls VE-cadherin localization and blood vessel stability. Nat Commun. 2015;6:6429.

37. Munger JS, Sheppard D. Cross talk among TGF-beta signaling pathways, integrins, and the extracellular matrix. Cold Spring Harb Perspect Biol. 2011;3(11):a005017.

38. Pulous FE, Petrich BG. Integrin-dependent regulation of the endothelial barrier. Tissue Barriers. 2019;7(4):1685844.

39. Mezu-Ndubuisi OJ, Maheshwari A. The role of integrins in inflammation and angiogenesis. Pediatr Res. 2021;89(7):1619–1626.

40. Leavesley DI, Schwartz MA, Rosenfeld M, Cheresh DA. Integrin beta 1- and beta 3-mediated endothelial cell migration is triggered through distinct signaling mechanisms. J Cell Biol. 1993;121(1):163–170.

41. Hutchings H, Ortega N, Plouet J. Extracellular matrix-bound vascular endothelial growth factor promotes endothelial cell adhesion, migration, and survival through integrin ligation. FASEB J. 2003;17(11):1520–1522.

42. Itkin T, Gur-Cohen S, Spencer JA, et al. Distinct bone marrow blood vessels differentially regulate haematopoiesis. Nature. 2016;532(7599):323–328.

43. Letteboer TG, Mager HJ, Snijder RJ, et al. Genotype-phenotype relationship for localization and age distribution of telangiectases in hereditary hemorrhagic telangiectasia. Am J Med Genet A. 2008;146A(21):2733–2739.

44. Bourdeau A, Dumont DJ, Letarte M. A murine model of hereditary hemorrhagic telangiectasia. J Clin Invest. 1999;104(10):1343–1351.

45. Hammill AM, Wusik K, Kasthuri RS. Hereditary hemorrhagic telangiectasia (HHT): a practical guide to management. Hematology Am Soc Hematol Educ Program. 2021;2021(1):469–477.

