## Supplementary material for "Endoglin regulates the integrity of the bone marrow vasculature": Materials and Methods

### Mice

All mice were housed under specific pathogen-free (SPF) conditions at the Experimental Centre of the Medical Theoretical Center (MTZ, Technical University of Dresden-University Hospital Carl-Gustav Carus, Dresden, Germany) or at the Max Planck Institute of Molecular Cell Biology and Genetics (MPI-CBG, Dresden, Germany). The *Eng*$^{f/f}$ mouse line has been described previously and contains *loxP* sites flanking exons five and six of the endoglin gene.[21] The Cdh5:CreERT2 mouse line was kindly provided by Ralf Adams (Münster, Germany)[22] and was used to generate Cdh5:CreERT2-*Eng*$^{f/+}$ and Cdh5:CreERT2-*Eng*$^{f/f}$. WT controls in all experiments were CreERT2-negative *Eng*$^{f/+}$ or *Eng*$^{f/f}$ littermates, respectively. Experiments were performed with male and female mice from eight weeks of age, and no significant differences between the genders of the same genotype were observed for any of the performed analyses within this study. Mice were genotyped using primers described in Supplemental Table 1 and knock-down efficiency confirmed via immunofluorescence of BM sections (Figure 1B). Breeding of all mouse lines and animal experiments were in accordance with the local guidelines on animal welfare and were approved by the Landesdirektion Sachsen, Germany.

### Tamoxifen treatment

Tamoxifen was dissolved in corn oil (10 mg/ml) at 37°C and injected intraperitoneally once daily for five consecutive days (200µl per injection). After the injections, mice were fed a Genobios tamoxifen-containing diet (code GEN16/T400-R) for the duration of the experiment.

### Blood analysis

Red blood cells (RBC), platelets (PLT) and WBCs, including monocytes, neutrophils, eosinophils and lymphocytes were measured in whole blood using a Sysmex automated blood cell counter (Sysmex 117 XE-5000) as described in our previous work.[23]

### Flow cytometry

Bone marrow cells were isolated by crushing femurs in PBS containing 2% FCS using a mortar and then filtering the suspension through a 40 µm filter. Following centrifugation, the resulting pellet was resuspended in ACK lysis buffer for 30-40 seconds to lyse red blood cells, and the lysis was subsequently stopped with PBS containing 2% FCS solution. After centrifuging the solution, cells were stained for specific cell markers using fluorophore-conjugated antibodies for 30 minutes at +4 °C (see Supplemental Table 2 for more information on antibodies). After washing, cells were suspended in PBS containing 2% FCS solution for FACS analysis performed on either LSRFortessa (Becton Dickinson) or FACSCanto II (Becton Dickinson), and cell numbers were counted on MACS quant (Miltenyi Biotec).

**Cell cycle analysis with Ki67**

Surface markers of BM cells were stained as described in the flow cytometry section. Next, cells were fixed and permeabilized using the eBioscience kit for permeabilization, according to the manufacturer's instructions (00-5523-00). Afterwards, cells were stained using the Ki67 kit, according to the manufacturer's instructions (11-5698-82). Finally, cells were centrifuged, and the pellet was resuspended in PBS containing 2% FCS solution with DAPI (1:10,000) for analysis as described in the flow cytometry section.

**Cell viability analysis with Annexin V**

Before staining for surface markers, cells were treated with 1mg/ml of Collagenase A dissolved in DMEM with 10% FCS for 30 minutes at 37°C with gentle shaking. The reaction was halted by adding a 2% FCS solution and centrifuging the cells. Subsequently, cells were stained with fluorophore-conjugated antibodies as described in the flow cytometry section. Following centrifugation, cells were also stained with Annexin V apoptosis detection kit (BD Biosciences, cat. 556547), following the manufacturer's instructions. Finally, cells were centrifuged, and the pellet was resuspended in a 2% FCS solution with DAPI (1:10,000) for analysis as described in the flow cytometry section.

**Total (including intra- and extracellular) integrin staining**

Before staining surface markers, cells were fixed and permeabilized with the eBioscience kit for permeabilization (00-5523-00), according to the manufacturer's instructions. Afterward, cells were stained with fluorophore-conjugated antibodies against specific cell markers and target integrins and analyzed as described in the flow cytometry section.

**Extracellular integrin staining**

Cells were obtained as described in the flow cytometry section. To maintain consistent fluorophore activity between permeabilized and non-permeabilized conditions, cells were fixed with the fixation buffer from the eBioscience kit (00-5523-00) before staining for surface markers. After washing and centrifuging, cells were stained with fluorophore-conjugated antibodies against specific cell markers and target integrins as described in the flow cytometry section. After washing and centrifuging, cells were permeabilized with the eBioscience kit for permeabilization (reference number 00-5523-00), according to the manufacturer's instructions. Following washing and centrifugation, cells were analyzed as described in the flow cytometry section.

**Next generation sequencing**

Flow cytometry and cell sorting were performed as described in the previous section, using Aria II (Becton Dickinson). [24] ECs from the BM were defined as $CD45^-$, $Ter119^-$ and $CD31^+$, $Sca1^+$. Cells were isolated directly into the lysis buffer of the RNeasy Plus Micro Kit. RNA was isolated according to the manufacturer's instructions, and SmartSeq2 sequencing was performed.

**Transcriptome Mapping**

Low quality nucleotides were removed utilizing the Illumina fastq filter (http://cancan.cshl.edu/labmembers/gordon/fastq_illumina_filter/). Reads were further subjected to adaptor trimming using cutadapt.[25] Alignment of the reads to the Mouse Genome was done using STAR Aligner using the parameters: "––runMode alignReads ––outSAMstrandField intronMotif––[2626] outSAMtype BAM SortedByCoordinate ––readFilesCommand zcat." [26] Mouse Genome version GRCm38 (release M12 GENCODE) was used for the alignment.

**Read Quantification**

Using the parameters: 'htseq-count -f bam -s reverse -m union -a 20', HTSeq-0.6.1p1[27] was used to count the reads that map to the genes in the aligned sample files. The GTF file (gencode.vM12.annotation.gtf) used for read quantification was downloaded from Gencode (https://www.gencodegenes.org/mouse/release_M12.html).

**Differential Expression Analysis**

Gene centric differential expression analysis was performed using DESeq2_1.8.1.[28] Also, the raw read counts for the genes across the samples were normalized using "rlog" command of DESeq2 and subsequently these values were used to render a PCA plot using ggplot2_1.0.1.[29] Heatmaps were generated using ComplexHeatmap package of R/Bioconductor.[30]

**Functional Analyses**

Pathway and functional analyses was performed using GSEA[31] and EGSEA.[32] GSEA/EGSEA were run using a normalized gene expression matrix against databases like Molecular Signatures Database (MSigDB), Reactome, KEGG and GO based repositories.

**Proteome profiler**

One femur was crushed in 200µl of PBS using a mortar, then centrifuged and the supernatant was collected and stored at -80°C until analysis. Three samples were pooled together, and the protein concentration was measured using Abcam's Pierce BCA protein assay kit (cat 23227). A total of 75 µg of protein were used for the assay, which was conducted according to manufacturer's instructions (R&D systems, cat. ARY015). The resulting membrane was imaged using Fusion Fx (PEQLAB) and the mean gray value of each individual spot was measured using FIJI (ImageJ distribution 1.53s) by adjusting the region of interest (ROI) around each spot. Mean gray values were normalized against the average of the mean gray values of the control spots provided on the membrane.

**Protein analysis**

Plasma was collected from whole blood and centrifuged at 3000 RPM for ten minutes at 4°C. Meso Scale Discovery (MSD, Rockville, Maryland) was used for quantitative determination of the cytokines (IL-1β, IL-2, IL-4, IL-6, IL-8, IL-10, IL-13, IFN-γ, and TNF-α) using 50 µl of plasma in the Proinflammatory Panel 1 (mouse) V-PLEX Kit and MSD plate reader (QuickPlex SQ 120). Cytokine concentrations were calculated by converting the measured MSD signal to pg/ml using a standard. All values below the "blank" samples were considered as zero.

**Bone isolation for histological analysis, bone structure analysis and histomorphometry**

Mouse femurs were isolated and incubated overnight with 4% PFA. Next, they were treated for 96 hours with Sigma-Aldrich's osteosoft (1017281000) at 37°C. Finally, they were embedded in GmbH's Tissue-Tek (4583), frozen on dry ice, and kept at -20°C until sectioning.

**Bone sectioning, staining, and microscopy image acquisition**

Seven µm mouse femur cryo-sections were blocked for one hour using Agilent's blocking solution (X090930-2). Next, sections were incubated overnight at 4°C with primary antibodies diluted in Agilent's antibody diluent (S302283-2) to detect target antigens. The following day, sections were incubated with fluorescently labeled secondary antibodies (see Supplemental

Table 2 for more information on antibodies). Images were acquired on an ApoTome II Colibri (Carl Zeiss, Jena, Germany) with a Zeiss Plan-Apochromat 10X/0.45. Ph1 objective and a Monochrome AxioCam 506 mono camera for a resulting 4.4053 pixels/µm image with a depth of 14 bits. Images were analyzed using either Zen software (Carl Zeiss, Jena, Germany) or Fiji (ImageJ distribution 1.53s) as described in the following sections.

**Vessel area mode**

After staining for endomucin and Eng and imaging of femur cryosections as described above, bone marrow images were divided into individual images using Fiji. Next, all blood vessels in each image were manually selected and individual areas were defined using FIJI's ROI group property. The peak of the distribution of all measured vessel areas per section (referred to as "mode") was determined as shown in Supplementary Figure 1B. For this, all vessel areas of the were plotted together, and the peak of the distribution was calculated using RStudio (version 1.2.1335).

**Nearest neighbor analysis for Eng$^-$ vessels**

Individual Eng-negative blood vessels were manually selected as described in the previous section. The centroid of each remaining ROI (describing an Eng$^-$ vessel) was defined in Fiji and the average nearest neighbor distance (ANND) was calculated for each BM section.

**Dextran injection and analysis**

Dextran 0.25mg conjugated with Texas Red diluted in 100µl of PBS was injected intravenously via the tail vein. After 10 minutes, mice were anesthetized with Ketamine/Xylazine and perfused with 20ml of PBS followed by 20ml 2% PFA. Both femurs were isolated and processed as described previously. After staining with endomucin and image acquisition, the images were divided into individual ROIs using Fiji. The dextran$^+$ area was identified and separated for each ROI using an automated procedure implemented in the IJ1 macro language. The automated procedure consisted of processing the image with a Gaussian blur of $\sigma$=2 pixels, followed by segmentation using an individual threshold calculated for each ROI using Fiji's moments method and increasing the calculated threshold by 200 (from a 14-bit image). To calculate the dextran$^+$ area per µm² for each mouse femur, we divided the total area of dextran+ regions in all regions of interest (ROI) by the total summed area of all ROIs for the same mouse.

**Micro-computed tomography**

We obtained the bone volume and total volume of mice's spines using the µCT50 system (Scanco Medical AG, Switzerland) as previously described.[33] Briefly, the fourth lumbar vertebra was scanned at an isotropic voxel size of 10.5 µm with an integration time of 200 ms, X-ray intensity of 140 µA, and energy of 70 kV (vivaCT40, Scanco Medical AG, Switzerland). Within vertebrae, 100 slices (50 above and 50 below the center) were contoured to evaluate the trabecular bone structure using predefined scripts from Scanco.

**Statistics**

All data are presented as mean ± SEM. Data (Cre$^-$ control versus Cre$^+$) were analyzed using a two-tailed Mann–Whitney U-test, or an unpaired t-test with Welch's correction as appropriate (after testing for normality with the F-test), unless otherwise stated in the text. All statistical analyses were performed using GraphPad Prism v7.02 or higher for Windows (GraphPad Software, La Jolla California USA, www.graphpad.com).

**Data Sharing Statement**

The RNAseq data is available in the Gene Expression Omnibus database (accession number **GSE271409**).

For original data, please contact
