## Supplementary material for "Endoglin regulates the integrity of the bone marrow vasculature": Suppl. Figures

Supplementary Figure 1

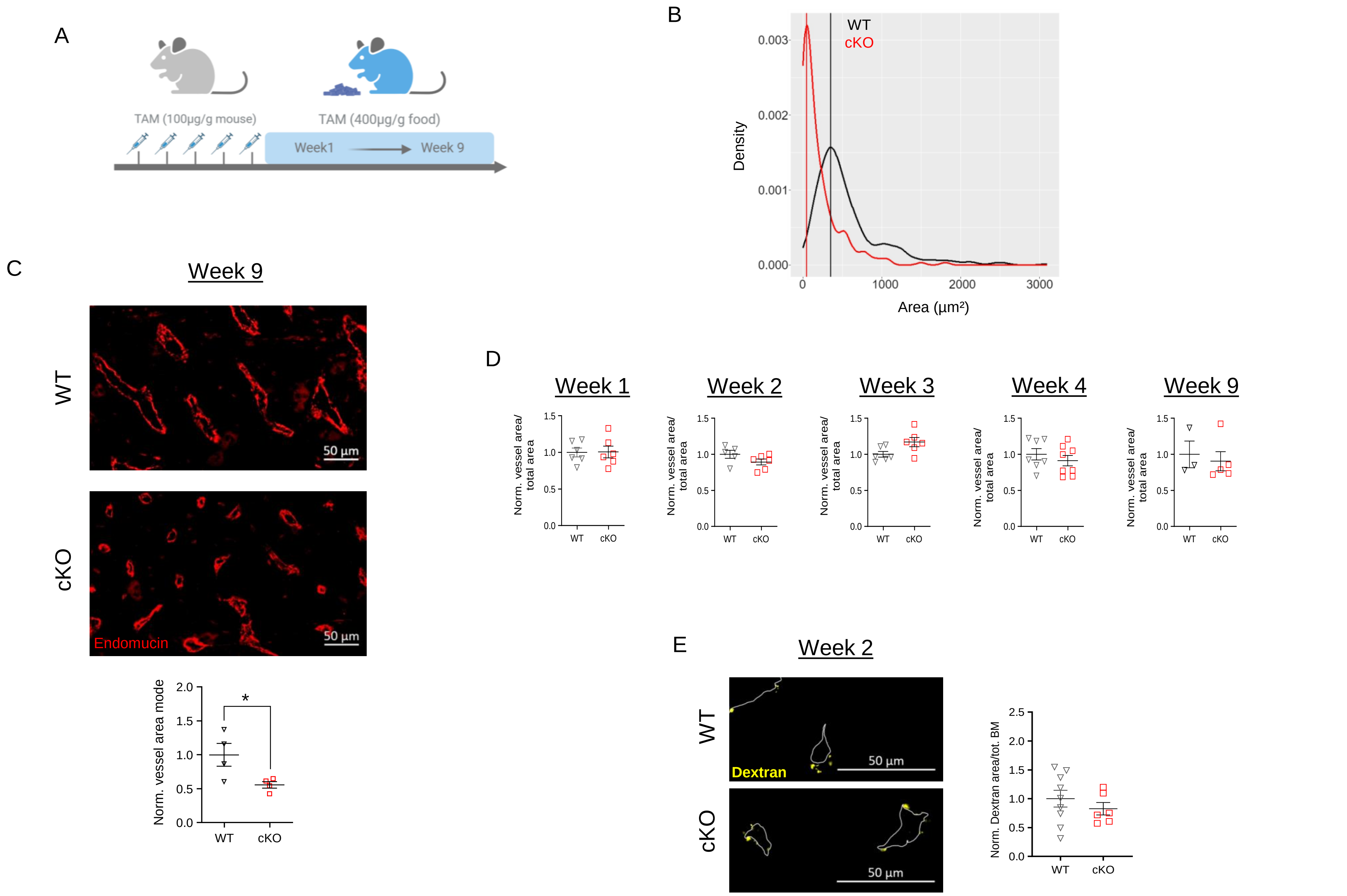

Supplementary Figure 2

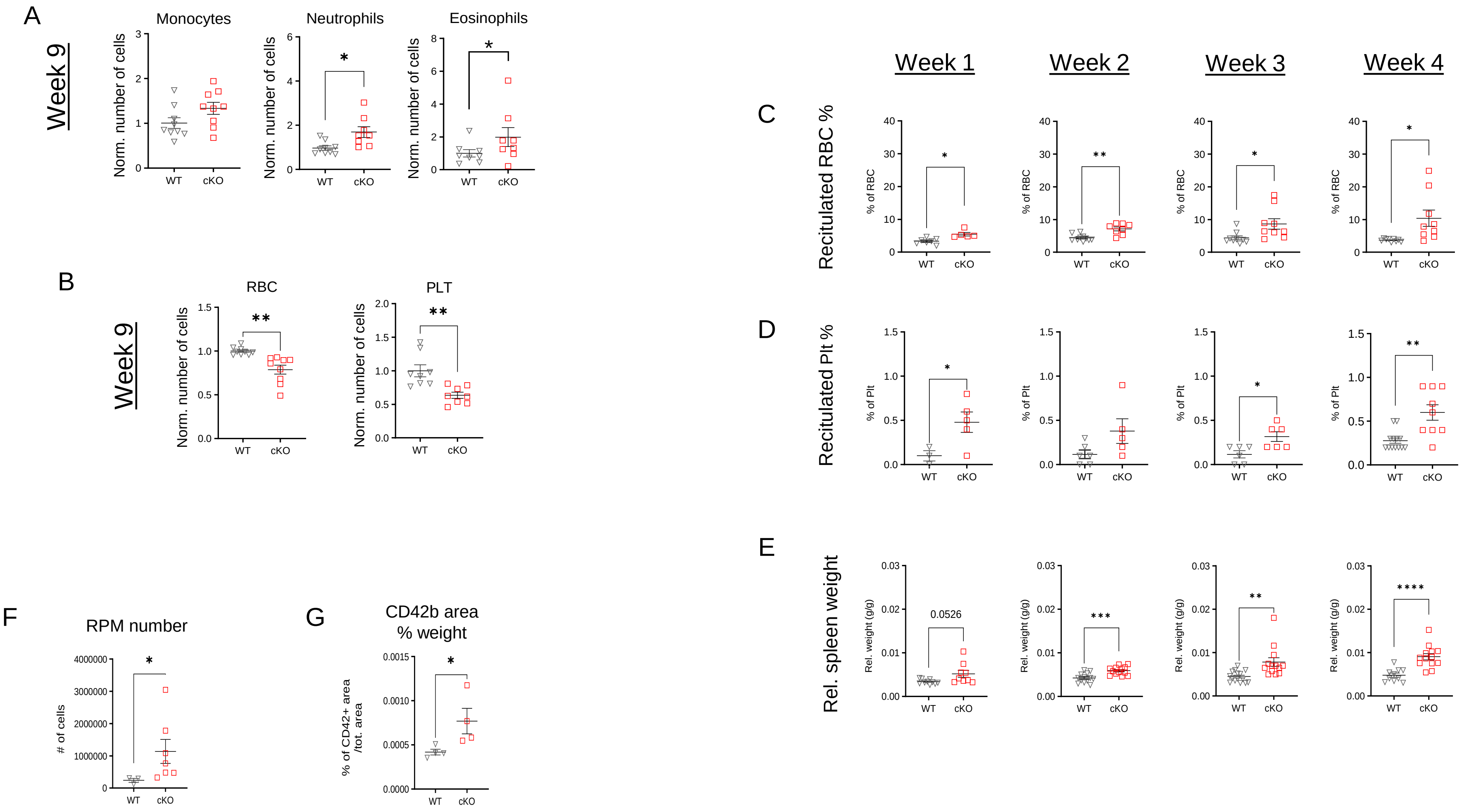

Supplementary figure 3

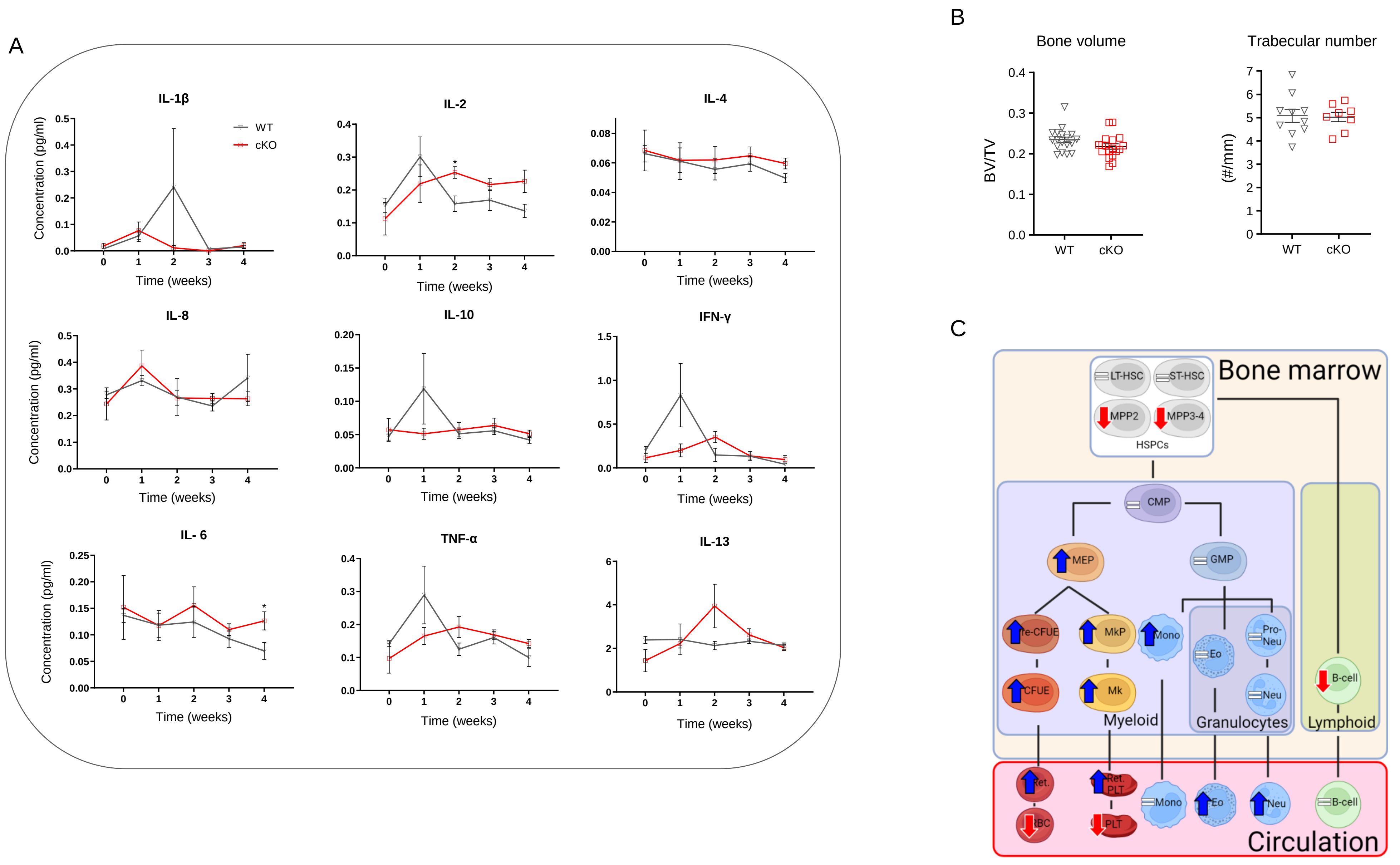

Supplementary Figure 4

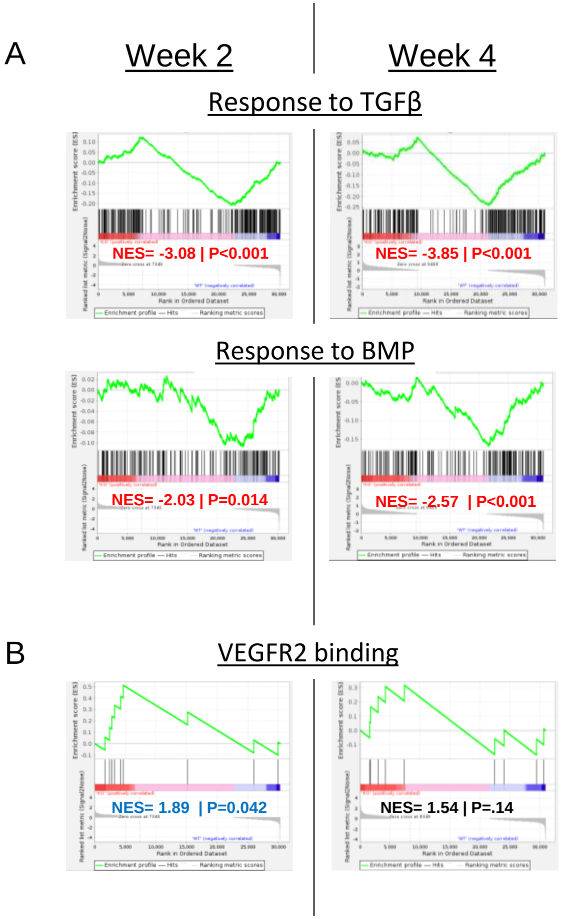

Supplementary Figure 5

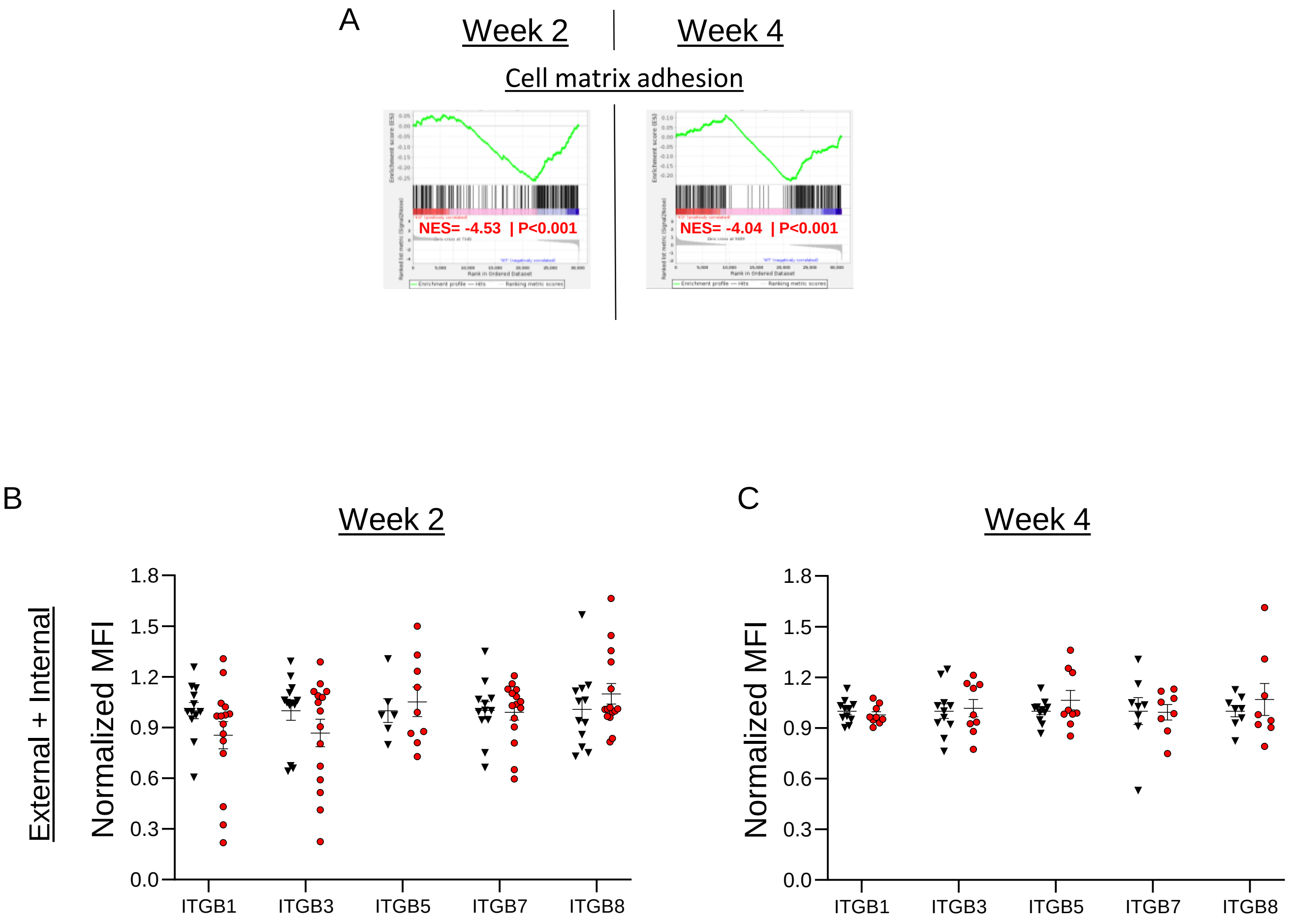

Supplementary Figure 6

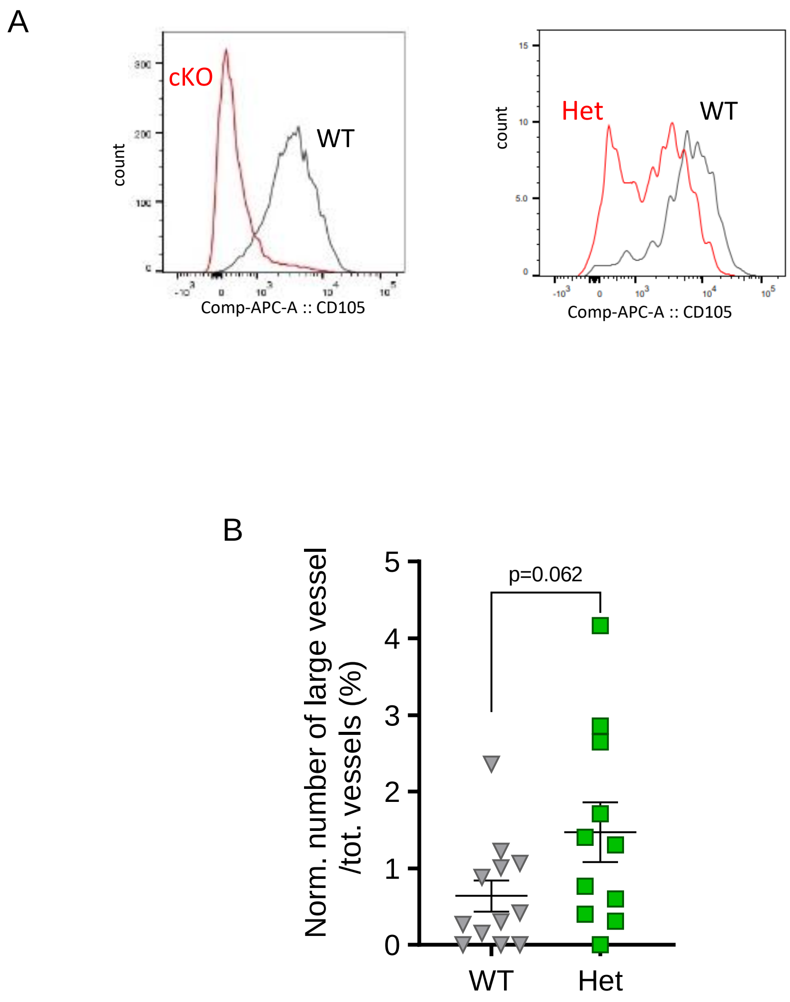

Supplementary Figure 7

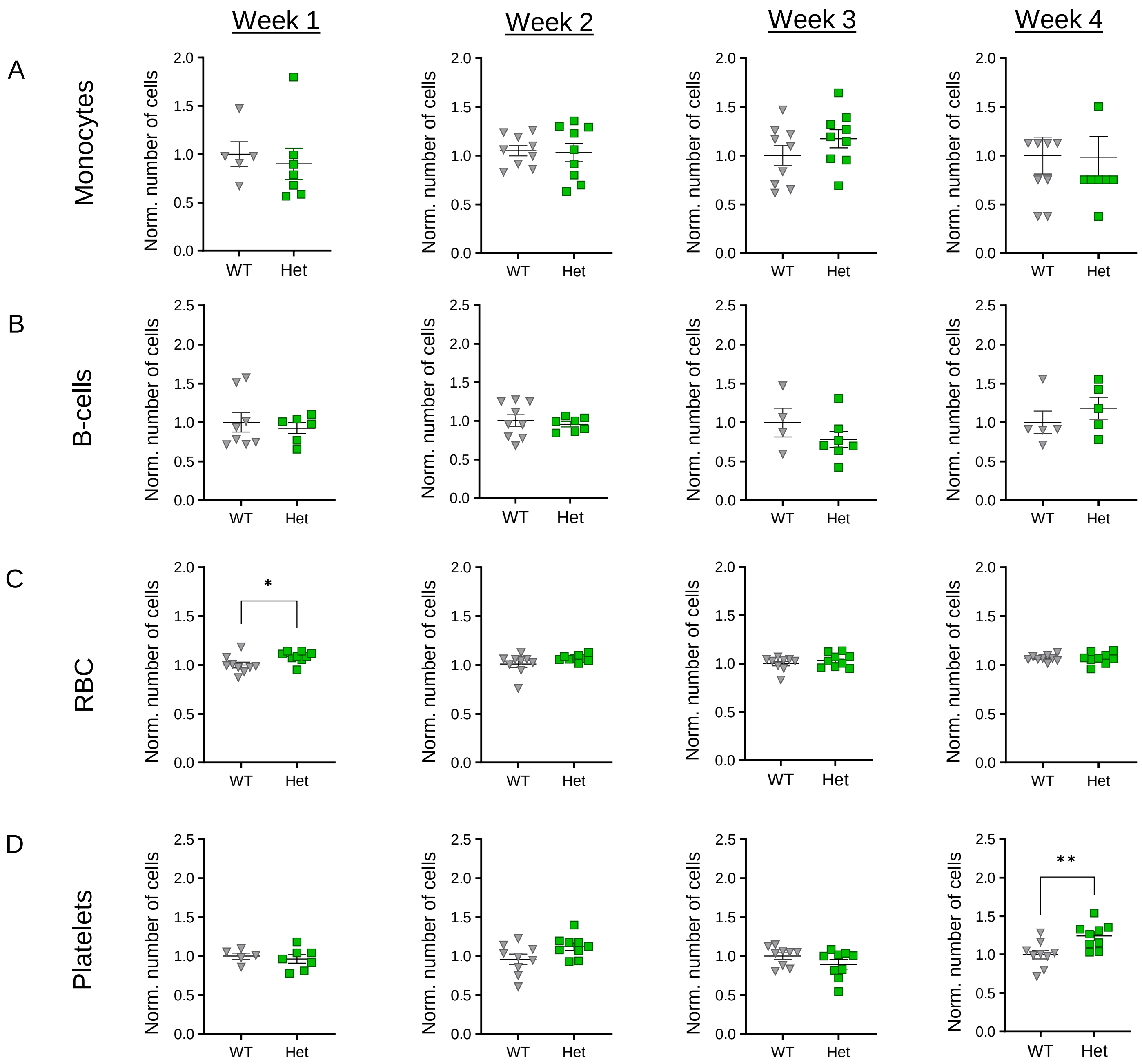

Supplementary figure 8

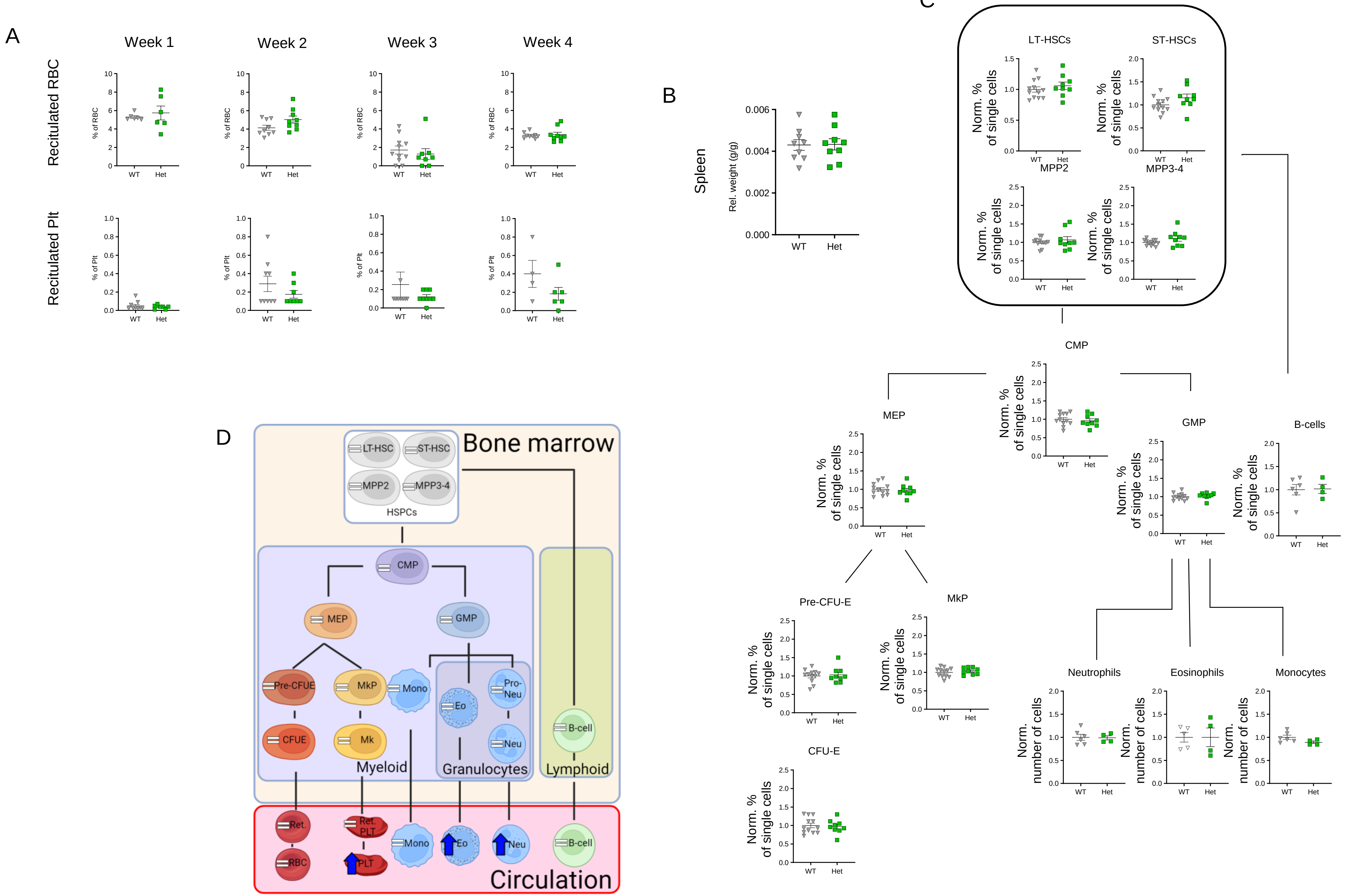
