## Supplementary material for "Endoglin regulates the integrity of the bone marrow vasculature": Suppl. Tables

**Supplemental Table 1. List of primers used to genotype mice**

| | |
|---|---|
| Endoglin flox 1 | 5' GGTCAGCCAGTCTAGCCAAG 3' |
| Endoglin flox 2 | 5' GTGGTTGCCATTCAAGTGTG 3' |
| Cre 1 | 5´ GCCTGCATTACCGGTCGATGCAACGA 3´ |
| Cre 2 | 5´ GTGGCAGATCGCGCGGCAACACCATT 3´ |

| **Supplemental Table 2. List of antibodies** | | | | |
|---|---|---|---|---|
| **Primary Antibodies** | | | | |
| **Antibody** | **Dilution** | **Host** | **Catalog number** | **Company** |
| CD3e bio | 1:1000 | Hamster | 13-0031-82 | ebioscience |
| CD19 bio | 1:500 | rat | 13-0193-81 | ebioscience |
| NK1.1 bio | 1:2000 | mouse | 13-5941-81 | ebioscience |
| Ter119 bio | 1:500 | rat | MA5-17819 | ebioscience |
| Ter119 PE-Cy7 | 1:200 | rat | 25-5921-81 | ebioscience |
| CD11b bio | 1:200 | rat | 13-0112-81 | ebioscience |
| GR-1 bio | 1:800 | rat | 13-5931-82 | ebioscience |
| B220 | 1:400 | rat | 13-0452-82 | ebioscience |
| CD16/32 Alexa fluor 700 | 1:50 | rat | 56-0161-82 | ebioscience |
| CD48 APC | 1:300 | Armenian hamster | 103411 | Biolegend |
| CD150 PE-Cy7 | 1:50 | rat | 115914 | Biolegend |
| cKit APC-efluor 780 | 1:600 | rat | 47-1171-80 | ebioscience |
| Sca1 PE-Cy5 | 1:100 | rat | 15-5981-82 | ebioscience |
| Sca1 Alexa fluor 700 | 1:200 | rat | 56-5981-80 | ebioscience |
| CD105 PE | 1:200 | rat | 12-1051-82 | ebioscience |

| | | | | |
|---|---|---|---|---|
| CD41 PerCP-eF710 | 1:400 | rat | 46-0411-82 | ebioscience |
| CD34 FITC | 1:50 | rat | 11-0341-85 | ebioscience |
| CD45 PE-Cy7 | 1:700 | rat | 25-0451-81 | ebioscience |
| CD31 APC | 1:600 | rat | 17-0311-80 | ebioscience |
| CD31 PE-e610 | 1:300 | rat | 61-0311-82 | ebioscience |
| CD62e PE | 1:200 | rat | sc-59766 PE | Santa Cruz |
| CD105 PE | 1:200 | rat | 12-1051-82 | ebioscience |
| CD29 APC-eFluor 780 | 1:200 | hamster | 47-0291-82 | Invitrogen |
| CD18 FITC | 1:100 | rat | 11-0181-82 | Invitrogen |
| CD61 APC | 1:100 | Armenian hamster | 17-0611-82 | Invitrogen |
| ITGB5 FITC | 1:100 | mouse | 11-0497-41 | Invitrogen |
| ITGB7 bio | 1:100 | rat | 13-5867-82 | Invitrogen |
| ITGB8 | 1:100 | rabbit | PA5-100843 | Invitrogen |
| Endomucin | 1:200 | goat | **PA5-47648** | Thermo Scientific |
| **Secondary Antibodies** | | | | |
| **Antibody** | **Dilution** | **Host** | **Catalog number** | **Company** |
| SA eF450 | 1:300 | | 48-4317-82 | ebioscience |
| Anti-Rabbit-IgG A647 | 1:500 | donkey | ab150075 | Abcam |

| | | | | |
|---|---|---|---|---|
| Anti-Goat-IgG<br>A647 | 1:350 | donkey | A-21447 | Abcam |
| Anti-Goat-IgG<br>A555 | 1:350 | donkey | ab150130 | Abcam |
| Anti-rabbit-IgG<br>A488 | 1:350 | donkey | A-21206 | Thermo Scientific |
